# ChipNMR: Hyperpolarized NMR for non-invasive metabolic flux analysis in perfused microfluidic chips

**DOI:** 10.1101/2025.05.20.654798

**Authors:** Thomas B. Wareham-Mathiassen, Juan D. Sánchez-Heredia, Ke-Chuan Wang, Cajsa R. Haupt, Magnus Karlsson, Alexander Jönsson, Martin Dufva, Roland Thuenauer, Pernille Rose Jensen

## Abstract

Dissolution dynamic nuclear polarization NMR Spectroscopy (dDNP-NMR) has become a transformative tool for metabolic studies by significantly enhancing signal sensitivity more than three orders of magnitude compared to traditional NMR. However, NMR detection probes are optimized for round narrow glass tubes typically 5 mm in diameter, which impose constraints on their utility for metabolic studies of adhernt cells. Here, we present a novel NMR probe head integrated with a custom microfluidic chip that facilitates real-time monitoring of hyperpolarized substrate conversion from adhernt cells. This system enables metabolic flux analysis in a controlled, in vitro environment, as demonstrated by tracking the conversion of [1-^13^C] pyruvate to [1-^13^C] lactate in HeLa cells over 48 hours. To the best of our knowledge, this is the first demonstration of cell metabolism from an adhering monolayer of mammalian cells in combination with hyperpolarized NMR. The custom microfluidic chip design is modular and adaptable allowing expansion to dual-chamber chips, demonstrating its potential in applications for more complex cellular environments, such as Organ-on-a-Chip systems.

## Introduction

The process of drug development is both resource-intensive and time-consuming, with high attrition rates often stemming from the limited predictive power of traditional preclinical models^1^. Historically, animal models have been the cornerstone of drug discovery, yet their translational relevance to human physiology remains suboptimal, leading to costly failures in clinical trials. This limitation underscores the necessity for more accurate and human-relevant *in vitro* models that better replicate the complexity of human disease^2^.

Conventional two-dimensional (2D) cell culture systems, while widely used, fail to recapitulate key aspects of the physiological microenvironment, including cell-cell interactions, extracellular matrix composition, and dynamic biochemical gradients^3^. Three-dimensional (3D) cultures provide improved structural and functional relevance, but they still lack precise control over perfusion, mechanical forces, and biochemical signaling present *in vivo*^4^. Organ-on-a-chip (OOC) technology addresses these limitations by integrating microfluidic systems that simulate organ-level functions, providing a physiologically relevant platform for studying cellular behaviors, disease mechanisms, and drug responses^5–11^.

Despite the advances in OOC platforms, a major challenge persists: the lack of methods to directly and continuously monitor metabolic activity within perfused systems in real time. Metabolic alterations often precede morphological and functional changes in cells, making metabolism a key early indicator of disease progression, therapeutic response, and cellular adaptation^12–15^. Current methods for metabolic assessment in OOCs predominantly rely on indirect measurements, such as luminescence-based sensors for glucose consumption, oxygen utilization, and pH changes^16–20^. For example, electrochemical glucose and lactate biosensors operate via enzyme-mediated conversion to hydrogen peroxide, which is then quantified^*21*^. Electrochemical biosensors, while offering higher specificity, are also susceptible to signal drift, enzyme degradation, and the need for periodic calibration^20,22,23^. Several studies highlight the value of embedding sensors directly within chips to enhance the accuracy of metabolic measurements^21,23–25^. While these methods provide valuable metabolic insights, they lack the direct detection capability of real-time metabolic fluxes.

Nuclear Magnetic Resonance (NMR) spectroscopy is a powerful analytical tool for metabolic studies due to its noninvasive nature and ability to provide quantitative insights into metabolic pathways. In particular, dissolution dynamic nuclear polarization (dDNP) NMR significantly enhances signal sensitivity, enabling real-time monitoring of metabolic processes in living systems^26^. The potential synergies between dDNP-NMR and microfluidic OOC systems have been recognized, yet practical implementations remain limited^27^. Previous work has demonstrated the feasibility of integrating ^1^H NMR with microfluidic devices, as well as hyperpolarized ^13^C detection of signal from parahydrogen-induced polarization (PHIP) ^28,29^. However, no platform to date has fully integrated dDNP-NMR with a perfused OOC system for longitudinal metabolic assessment.

In this study, we present a novel NMR probe integrated with a custom-designed microfluidic chip, enabling real-time metabolic analysis in perfused cell cultures. This platform allows for monitoring of hyperpolarized substrate conversion within a physiologically relevant microenvironment. We demonstrate the functionality of this system through longitudinal metabolic measurements of HeLa cells, tracking the conversion of hyperpolarized [1-^13^C] pyruvate to [1-^13^C] lactate over 48 hours. Additionally, comparable kinetics are obtained in a more advanced chip with two chambers. This work establishes a foundation for combining OOC technology with noninvasive metabolic monitoring, providing deeper insights into cellular metabolism with broad implications for drug development, personalized medicine, and infection research.

### Experimental Section

All chemicals were purchased from Sigma-Aldrich unless otherwise stated.

### RF Coil Design and Creation

The NMR probe head design includes two separate RF coils, and a support structure made of 3D printed resin (Formlabs, USA). It is made for a 400 MHz NMR spectrometer (Bruker Avance NEO). Coil 1 is a saddle-shaped coil, used exclusively for the ^13^C nucleus. The two rectangular loops are separated 4.5 mm (with the width of the chip being 4 mm), in order to maximize the filling factor. The coil is made of silver wire, with a wire diameter of 1 mm. This coil geometry provides an inductance of approximately 130 nH at the frequency of ^13^C (100.55 MHz). The measured unloaded Q-factor is 78. The loaded Q-factor (when loaded with a chip filled with 40 mM phosphate buffer) is 64. Coil 2 is double tuned, to cover the ^1^H and ^2^H nuclei. The coil has two opposed copper plates, acting similarly to a microstrip line. The separation between the two plates is 12 mm. The coil is made of copper and provides an inductance of approximately 49 nH at the ^2^H frequency, and 52 nH at the ^1^H frequency. An illustration of the full probe setup is shown in Figure 1.

**Figure 1:**
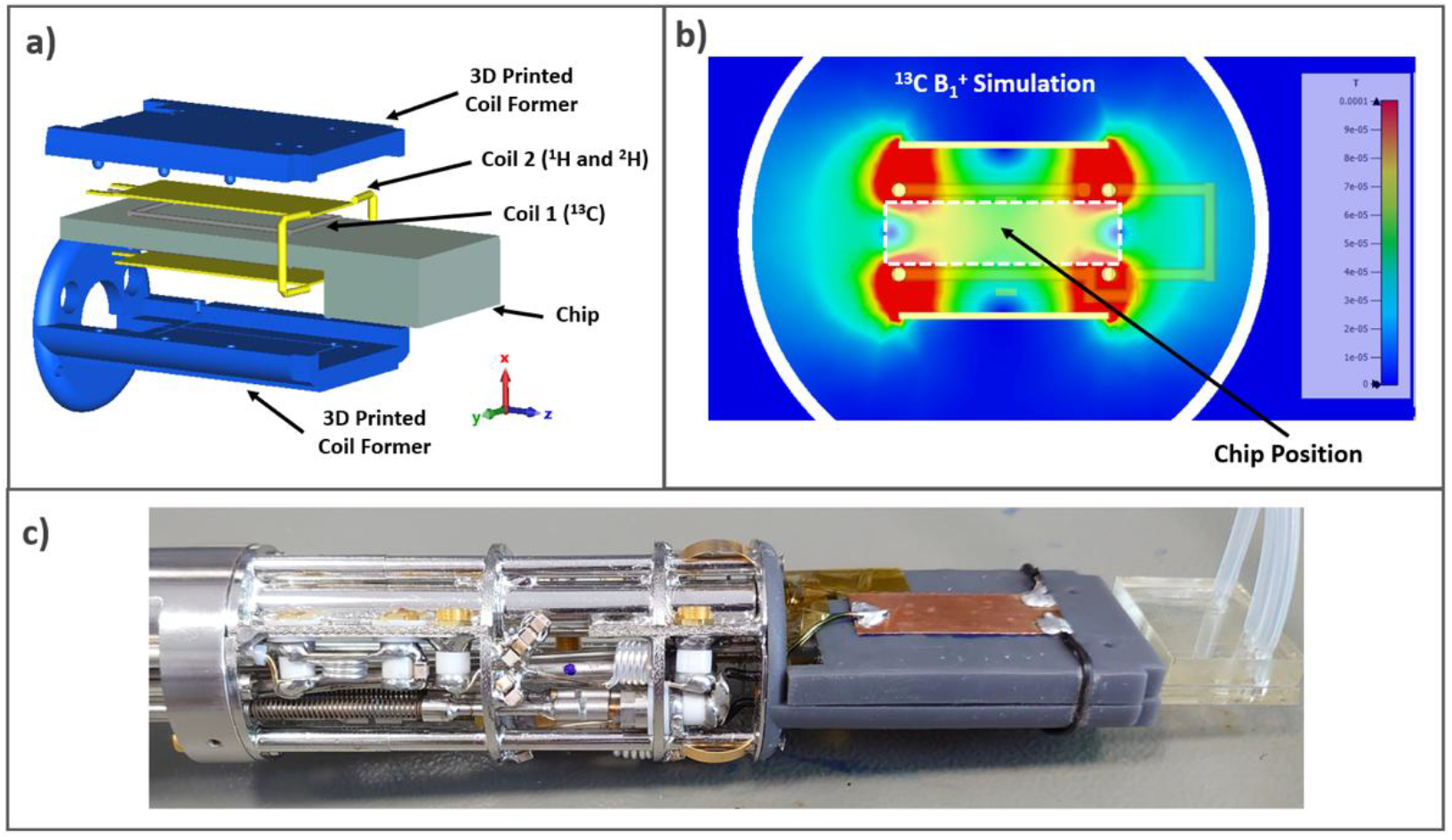
RF Coil design. (A) Exploded view of the NMR probe head, showing the two RF coils, the 3D-printed support, and the microfluidic chip. (B) Cross-section (Z = 0 plane) of the simulated B_1_^+^ field distribution generated by Coil 1 at the ^13^C frequency (per 1 W accepted power). (C) Photograph of the fabricated NMR probe with the chip inserted.

### Chip fabrication

The fluidic chip was fabricated using the following materials and procedures (Figure 2). Depending on the experimental design, either a 1-chamber or 2-chamber fluidic chip was fabricated. A 1 mm sheet of PETG (Exolon group, Vivak®) was covered with a 0.75 mm sheet of SEBS (Eden-Microfluidics, Flexym™), and a 0.3 mm sheet of PETG was covered with a 0.25 mm sheet of SEBS. For the 1-chamber chip, this bottom composite sheet was left uncut. These composite sheets were cut using a FLUX Beambox™ CO_2_ laser operated through Beam Studios™ software. The laser’s intensity, power, and speed were optimized to cut through the SEBS layer without penetrating the PETG layer, allowing for precise removal of the channel shape with tweezers. Subsequently, the outer shape of the chip was cut from the composite sheets and the SEBS side of a composite sheet was adhered to a sheet of track-etched polycarbonate (ipCELLCULTURE™ Track-Etched Membrane Filter with an 8 μm pore diameter, a pore density 1×10^5^/cm^2^ pore density, a thickness of 18 μm, and a porosity of 5%). The membrane was trimmed to fit using a scalpel, and the other composite sheet was pressed with its SEBS side on the opposite side of the membrane, forming a sandwich configuration as shown in Figure 2A. The assembled chip was then pressed at 2 Pa and 60°C for 1 hour using a hydraulic press. A top piece with four mini Luer injection ports was designed using SolidWorks and fabricated from BioMed Clear Resin on a Formlabs Form 3B printer with default settings and a layer height of 25 μm (Figure 2B). Post-printing, the inlets were washed with isopropanol and cured in a UV box at 70°C for 20 minutes. After curing, the supports were trimmed, autoclaved in 1 L of demineralized water at 121°C, and affixed to the chip using generic superglue

**Figure 2:**
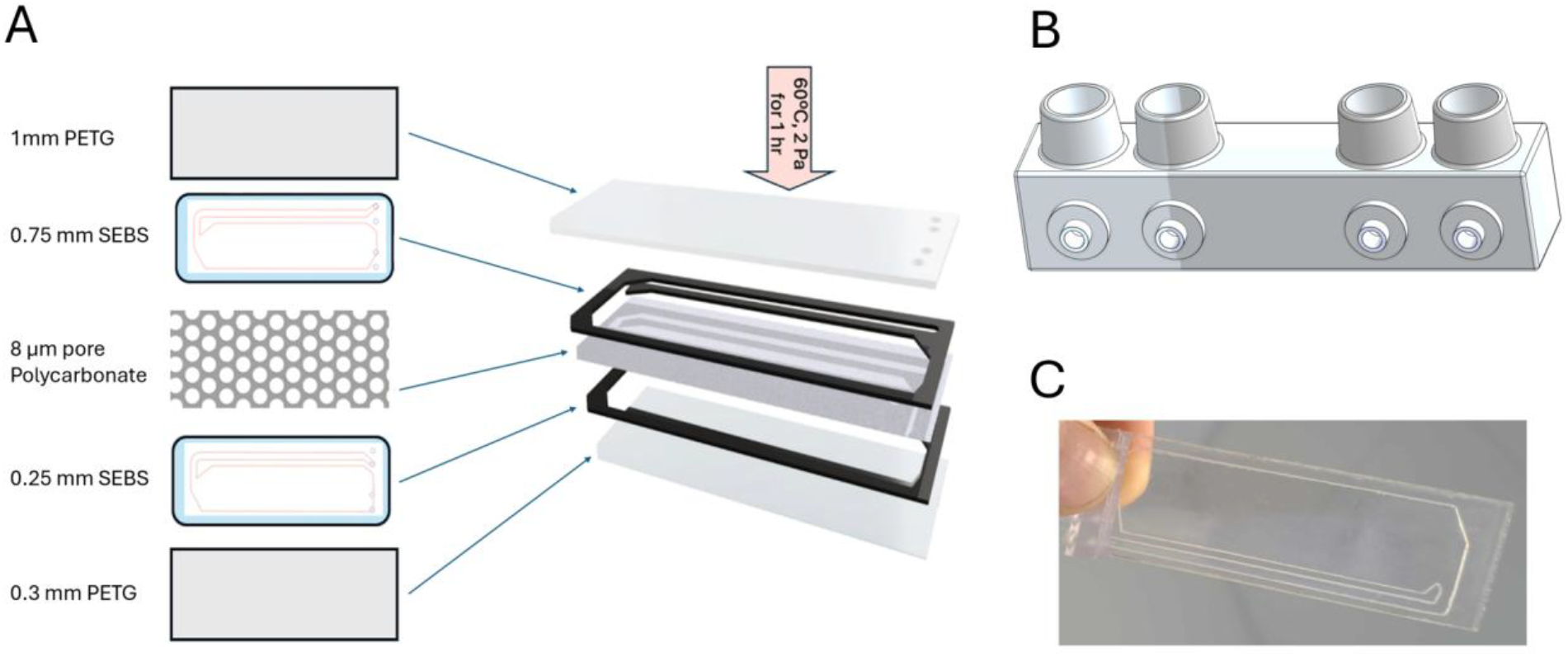
Fluidic Chip Production and Assembly: (A) Visual representation of the chip components and their assembly process. The stacked components are shown in order from top to bottom: 1 mm PETG, 0.75 mm SEBS, track-etched polycarbonate membrane (8 μm pore size, 18 μm thickness, 5% porosity), 0.25 mm SEBS, and 0.3 mm PETG. (B) 3D-printed inlets/injection ports, designed using SolidWorks, shown with female mini Luer connectors made from BioMed Clear Resin. (C) Fully assembled microfluidic chip.

### Chip Sterilization and Treatment

To ensure minimal leeching from the adhesives, the chip was injected with sterile H_2_O and placed in a water bath for 24 hours. Silicone tubes were attached to the chip with minileur connectors ChipShop product number 100000094), and 70% ETOH was perfused for 1 hour at 500μl/min using a peristaltic pump, the setup of which is outlined in previous work^30^. Following that, sterile water was flushed into the entire system for an additional 2 hours. Prior to seeding, the chips were coated with a solution containing 300 μg/mL Collagen I solution (Rat Tail Collagen Type I, Gibco) in DMEM (Biowest L0103), and placed at 37 °C for 1 hr.

### Cell Culturing and Chip Seeding

The HeLa human cervical cancer cell line (ATCC CCL-2, American Type Culture Collection, Manassas, VA) was routinely cultured in DMEM (Biowest L0103), which contains 4.5 g/L d-glucose, stable l-glutamine, and sodium pyruvate. The medium was supplemented with 10% fetal bovine serum (FBS; Biowest S1810) and 1% penicillin/streptomycin (Biowest L0022). Cells were maintained at 37°C in a humidified 5% CO_2_ incubator.

The cell medium was refreshed 2 to 3 times per week, and the cells were cultured until they reached 80% confluence, at which point they were harvested using a 0.25% Trypsin-EDTA solution (Biowest X0930). For the dDNP-NMR experiments, 1 million HeLa cells were seeded 4–5 days prior to the experiment in a T175 flask. On the day of seeding the chips the cells were harvested by trypsinization, followed by centrifugation and washing in DMEM containing 10 % FBS. Subsequently, the cells were washed in PBS with Ca2^+^/Mg2^+^ (HyClone, SH30264.01) and resuspended in DMEM + 10% FBS with to a final concentration of 4 × 10^6^ cells/mL.

The membrane surface area of each chip was 13 cm^2^, and chips were seeded at a density of 2.5 × 10^5^ cells/cm^2^, resulting in a total of 3 × 10^6^ cells per chip. Homogenized cell suspensions were injected into a vertically placed chip via the narrow channel using a 2 mL syringe fitted with a male mini Luer tip. Following seeding, the chips were incubated at 37°C in a humidified 5% CO_2_ atmosphere for three hours to allow cell adhesion, which was confirmed via visual inspection. Once adhesion was confirmed, a continuous flow of DMEM at 200 μL/min was initiated and maintained until the dDNP-NMR experiment commenced at 3, 24, or 48 hours.

### Cell Staining and Microscopy

The chips were washed twice with approximately 800 μL of PBS, then stained with a solution of 2 μg/mL Calcein AM (Invitrogen™ C1430) for 30 minutes at 37°C, protected from light. Following staining, the chips were washed again with PBS to remove excess dye. Fluorescence microscopy was performed using a Leica DMI3000 B microscope system (Leica, Wetzlar, Germany) with an excitation wavelength of 485 nm and an emission wavelength of 530 nm to visualize live cells. The number of fluorescent cells was used to quantify viable cells in each chip. Cell images were analyzed using ImageJ software (NIH, United States).

### dDNP-NMR Experiments

Substrate stock solutions were prepared using [1-^13^C] pyruvic acid (Sigma-Aldrich), doped with 17 mM trityl radical AH111501 (GE Healthcare) and 1.5 mM Gadoteridol (Bracco Imaging). Approximately 4.5 mg of the pyruvic acid stock solution was hyperpolarized using a SpinAligner 6.7 T polarizer (PolarizeTM) for ∼1 hour until equilibrium polarization was achieved. The sample was rapidly dissolved in 5 mL of phosphate buffer (pH 7.4, 40 mM), with 5 μL of 10 M NaOH added to stabilize the pH, maintaining a final temperature of ∼310 °K. The dissolved pyruvate concentration was ∼10 mM, with ∼25% liquid-state polarization. The solution was collected in a 50 ml receiver, and 3 mL was drawn into a 5 mL syringe, and 1.5 mL was injected into the chip sitting in the NMR spectrometer.

At designated time points when the pyruvate sample was fully polarized, a chip was fitted with an injection line of PEEK tubing, 1/32″ ID 0.5 mm (Microlab) and a silicon tube as outlet ID 1mm (Hounisen). Then placed into the NMR probe and inserted into a Bruker 400 MHz spectrometer, within 5 min.

^13^C NMR spectra were acquired over 256 s as a series of 128 1D ^13^C spectra each using one 10° pulse angle and a 2 s delay between pulses. The data acquisition was started manually upon start of the injection of hyperpolarized pyruvate into the chip.

### Cell Counting

Following the dDNP-NMR experiment, the chip was immediately removed from the spectrometer, cells were washed twice with PBS, followed by trypsinization for 10 minutes. The outflow was mixed with equal parts DMEM+10% FBS to neutralize the trypsin and spun down at 300 G before resuspending in 10 mL PBS. Cells were counted using an EVE™ Automated Cell Counter (nanoEntek) with 0.4% trypan blue stain.

### Data Analysis

The NMR data analysis was performed using MestReNova software (Mestrelab Research). After phase and baseline correction, the signals of the substrate ([1–^13^C] pyruvate) and the product ([1–^13^C] lactate) in the time series were integrated. A Python-based model was then applied to the integrals using a numerical solution of the ordinary differential equations (ODEs) that describe the forward reaction rate (k). In this model, the pulse length (pw) were fixed, while the T_1_ values (T1_S_, T1_P_) were allowed to vary. The ODEs used in the model were as follows:

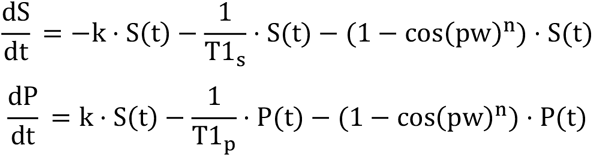

As it took eight seconds to fill up the chip with hyperpolarized pyruvate, the time curves for pyruvate were extrapolated back to the beginning of the injection to ensure an exponentially decreasing substrate curve.

### Statistical analysis

All statistical analysis was performed in GraphPad prism. p values were calculated with one-way ANOVA and values less than 0.05 were considered statistically significant.

## Results and Discussion

### NMR Coil Design and Functionality

The NMR probe head was designed to accommodate a flat microfluidic chip with a surface area as large as possible in order to allow for more than one million adhernt cells. The microfluidic chips were fabricated, leak-tested, and validated before experimentation. The corresponding chip included Luer-compatible injection ports, enabling the injection of hyperpolarized substrate while mounted in the probe. To validate the probe’s functionality, hyperpolarized [1-^13^C] pyruvate was injected into an empty chip, and the decay profile due to T1 relaxation was monitored by acquiring a time series of 1D ^13^C NMR spectra using a 10° pulse angle and a 2 s delay between pulses. A mono exponential T1 curve was observed demonstrating homogeneous filling of the chip (Figure 3A). The area under the curve (AUC), for three repetitions was 24 ± 0.6, with a coefficient of variation (CV) of 2.5%, demonstrating high reproducibility of the chip system. The signal-to-noise ratio (SNR) of the highest pyruvate peak spectrum was approximately 5000, confirming high sensitivity of the coil.

**Figure 3:**
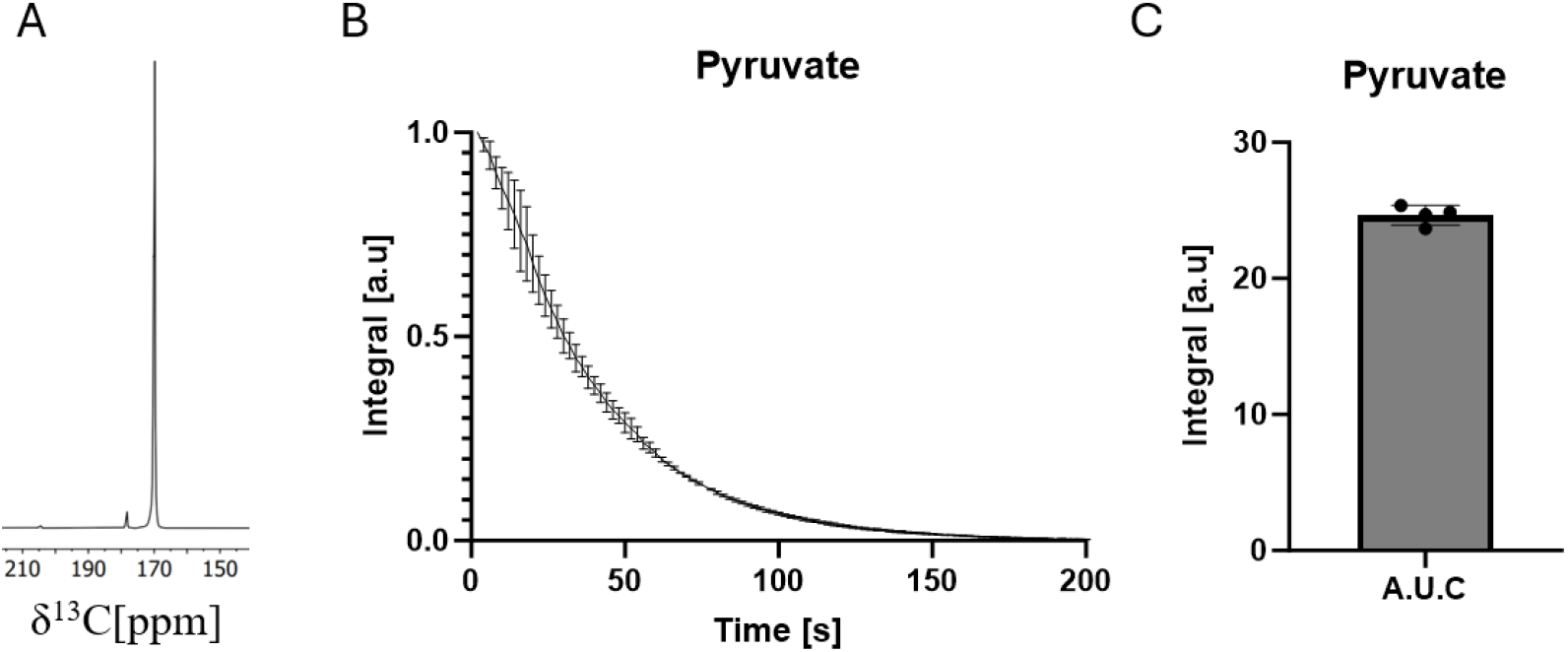
Performance Evaluation. A) Single pyruvate spectrum, SNR of the highest pyruvate peak was 5000, B) Substrate integral over time (n = 3) and C) comparative substrate integrals (AUC: 24 ± 0.6) following injection of 10 mM hyperpolarized [1-^13^C] pyruvate, CV for the AUC was 2.5%.

The custom-built NMR probe head, specifically engineered for microfluidic applications, demonstrated high reproducibility in the injection profile of the hyperpolarized substrate, achieving a coefficient of variation (CV) of 2.5% (Figure 3). Compared to conventional 5 mm NMR probes, which are generally incompatible with adherent cell cultures, our probe provides a tailored solution for microfluidic environments that support adherent cell cultures. This capability enables real-time monitoring of dynamic biological processes, offering a robust platform for advanced metabolic tracking and analysis. Unlike earlier NMR compatible microfluidic devices^31,32^, our setup has compromised on magnetic field homogeneity over the sample in exchange for larger cell surface area. This complementary strategy achieves a good balance between cell number, ease of use, and SNR. The large surface area of the chip allows the system to be operated via a simple peristaltic pump, in contrast to the more complex fluidic systems required in Southampton’s 2 μL microfluidic chip^28,33^. While this sacrifice in B1 homogeneity, leading to decreased peak separation ability, is not considered a significant issue for X-nuclei, it may present a bigger problem for ^1^H spectroscopy.

### Microfluidic Chip as a Cell Culturing Device

A custom microfluidic chip was developed in-house, specifically designed to optimize cell density within the NMR coil’s detection range. The chip was made transparent to allow for easy microscopic observation and to support fluorescence staining. Additionally, the chip was engineered to be biocompatible, flow-compatible, leak-proof, and resistant to material leaching, ensuring a stable and reliable environment for cell culture. Depending on the experimental design, either a 1-chamber or 2-chamber fluidic chip was used. Both configurations were equipped with female mini Luer injection ports on one end, allowing them to slide seamlessly into the dDNP-NMR probe without obstruction. The inlets were designed with protruding channels that extended into the chip, acting as a physical barrier to prevent affixing glue from entering the flow path.

To assess the chip’s ability to sustain cell culture, HeLa cell morphology and proliferation were monitored over 48 hours of culture. A total of 3 × 10^6^ HeLa cells were seeded onto the 1-chamber chips and incubated at 37°C with 5% CO_2_. After a 3-hour adhesion period, medium perfusion was initiated at a flow rate of 200 μL/min using a peristaltic pump. When necessary, flow was temporarily halted, and the chips were examined under a bright-field microscope before being returned to the incubator. At designated time points, cells were stained with the fluorescence stain Calcein AM for live cell imaging or detached using trypsin and counted with a cell counter employing trypan blue exclusion. Throughout the experiment, cells remained stably adhered to the chip’s membrane, displaying a highly confluent cell layer (Figure 4A) throughout the whole chip. Over 48 hours, the cell count doubled, further supporting the chip’s ability to sustain long-term perfused culture (Figure 4B). These results demonstrate the chip’s suitability for maintaining cell viability and proliferation under continuous perfusion conditions.

**Figure 4:**
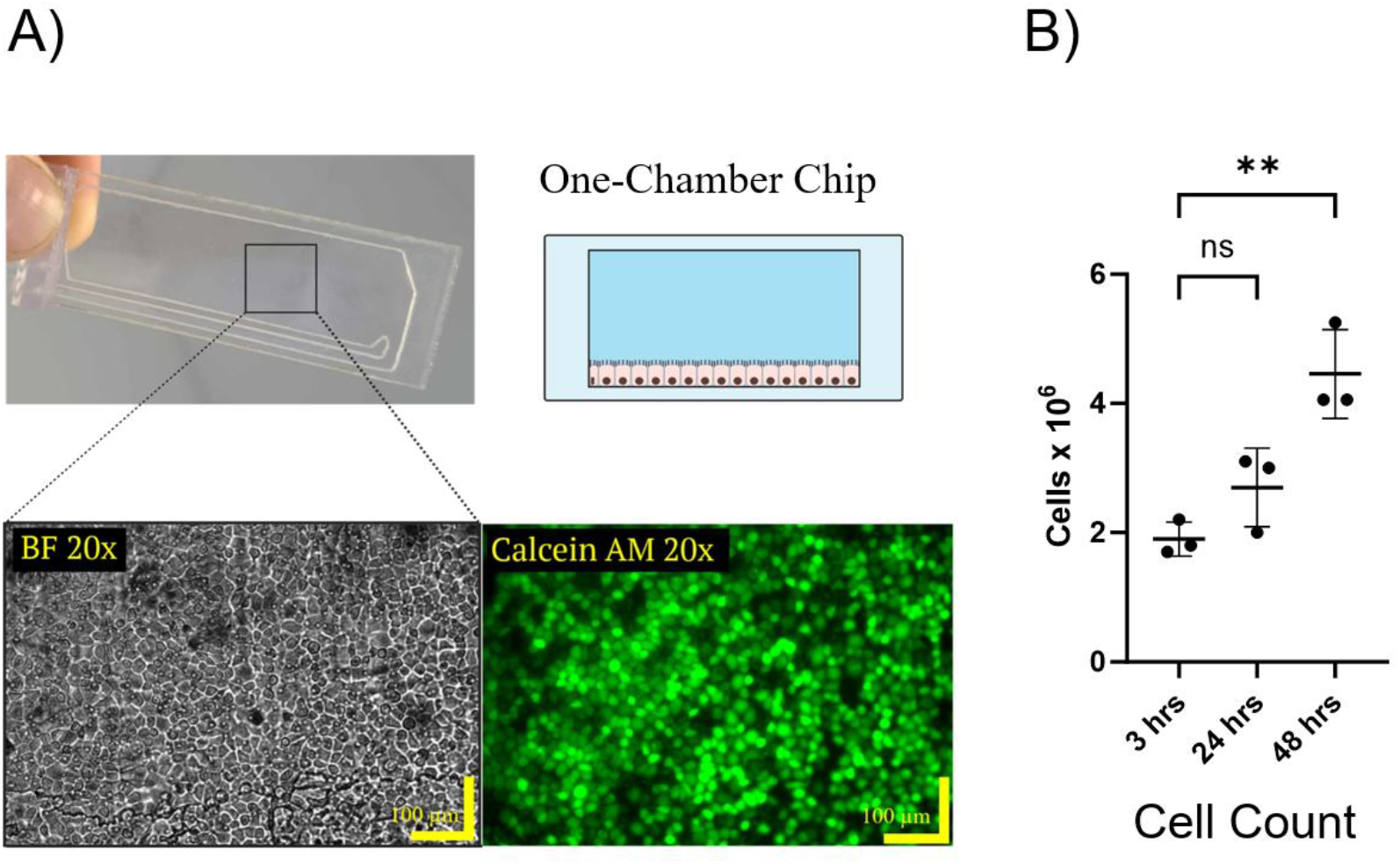
One-Chamber Chip Cell Growth. (A) One-chamber chip seeded with 3.5 × 10^6^ HeLa cells, cultured for 3 hours, and imaged with bright-field microscopy (bottom left, BF 20×) and fluorescence microscopy of cells stained with Calcein AM (bottom right, Calcein AM 20×). Scale bar: 100 μm. (D) Cell counts in chips after injection at 3, 24, and 48 hours. The mean ± SD cell counts (in millions) were 1.90 ± 0.26 at 3 hours, 2.70 ± 0.61 at 24 hours, and 4.46 ± 0.69 at 48 hours. Statistical analysis (one-way ANOVA) showed a significant difference (p < 0.01) in cell count between 3 and 48 hours. N = 3 for each time point.

### Detection of Metabolic Activity via dDNP-NMR

After validating the probe’s signal detection capabilities and confirming the microfluidic chip’s effectiveness as a cell culture platform, we assessed its ability to monitor metabolic activity in real time. Hyperpolarized [1-^13^C] pyruvate was injected into the mounted chip laden with HeLa cells, achieving a final concentration of 10 mM [1-^13^C] pyruvate. Metabolic conversion was tracked by monitoring lactate production from hyperpolarized [1-^13^C] pyruvate at 3, 24, and 48 hours after seeding into the chip. Figure 5A shows a representative spectrum from a time series 15 s after start of the injection where lactate has its maximum. The SNR of the highest lactate peak in this spectrum was approximately 50. The full dynamic curves showed a steady increase in metabolic activity over time (Figure 4B). The forward rate constants (k) were fitted to a set of differential equations using the integrals of the dynamic curves normalized to the initial pyruvate signal in each experiment (Figure 5B). The normalized rate constants were 4.12 ± 0.53 × 10^−4^ s^−1^ at 3 hours, 3.76 ± 0.64 × 10^−4^ s^−1^ at 24 hours, and 4.78 ± 0.53 × 10^−4^ s^−1^ at 48 hours. The corresponding coefficient of variation (CV) values were 13% at 3 hours, 17% at 24 hours, and 11% at 48 hours, with an average CV of 14%, indicating moderate variability in metabolic activity over the 48 hour period. Normalizing the rate constants to the increasing cell number between 3 and 48 hours resulted in the same turnover per cell throughout the experiment underscoring the stability of the system performance regarding metabolic flux determination (Figure 5C).

**Figure 5:**
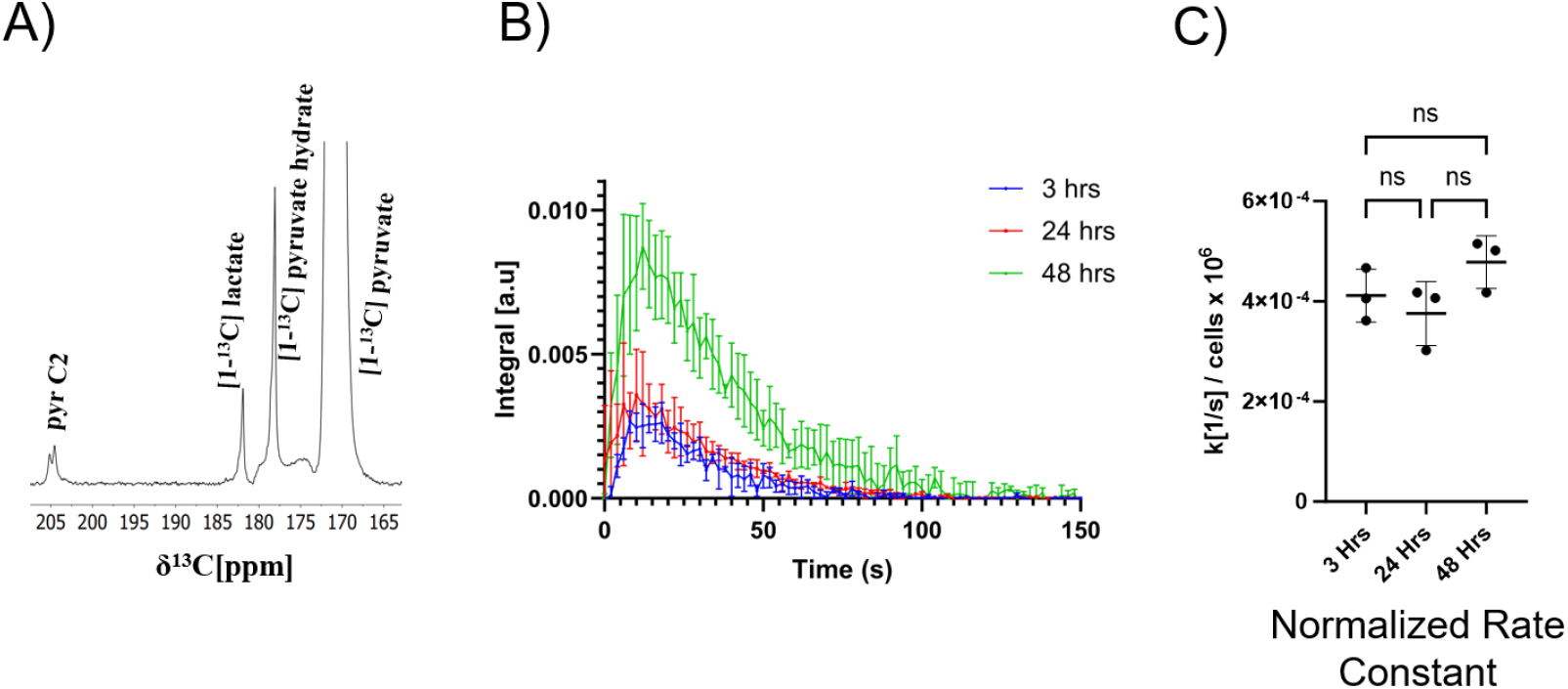
One chamber chip hyperpolarized ^13^C NMR. (A) Spectrum with the highest lactate signal (spectrum 9), showing pyruvate C2 at 208 ppm, lactate C1 at 184 ppm, pyruvate hydrate C1 at 178 ppm, and pyruvate C1 at 171 ppm.(SNR of highest lactate peak was approx. 50) (B) Time-series spectra displaying the production of [1-^13^C] lactate in one-chamber chips at 3, 24, and 48 hours post-seeding, following the injection of 10 mM hyperpolarized [1-^13^C] pyruvate. (C) Rate constants normalized to cell counts, yielding mean ± SD values (s^−1^) of (4.12 ± 0.53) × 10^−4^ at 3 hours, (3.76 ± 0.64) × 10^−4^ at 24 hours, and (4.78 ± 0.53) × 10^−4^ at 48 hours. Statistical analysis (one-way ANOVA) showed no significant difference (NS) between normalized rate constants. N = 3 for each time point.

An increasing total rate constant was observed as total cell numbers increased (Figure S1), whereas the rate constant normalized to cell number was stable (Figure 5). This observation shows the stability of the system where circulating medium removes dead cells. It is not possible to directly compare if the adherent cells have higher turnover compared to harvested cells because the suspended cells would be washed out of the chip upon injection of the hyperpolarized substrate. Nevertheless, in previous studies, we calculated the rate constants of pyruvate-to-lactate conversion in 2 million HeLa cells to approximately k = 2 × 10^−4^ s^−1^ (equivalent to 1 × 10^−4^ s^−1^ per million cells^34^. Our current results show comparable normalized rate constants (4 × 10^−4^ s^−1^ per million cells) consistent with the previous study. While the experimental setups were not identical, these results align with the idea that detaching cells can disrupt their metabolic processes^35–39^, and that measuring metabolism in adherent cells provides more physiologically relevant data. This corresponds with literature findings that show increased metabolic activity when conducting dDNP-NMR analysis on adherent cell culture compared with detached cells^40^.

High-quality flux data could be obtained on 2 × 10^6^ adherent cells. This sensitivity is comparable to data obtained from standard 5 mm probes on suspensions of cancer cells, where typically 2-20 × 10^6^ cells are used^34,41–44^. To the best of our knowledge, this is the first demonstration of cell metabolism from an adherent monolayer of mammalian cells in combination with hyperpolarized NMR. The CV of our metabolic measurements over the 48 hours was 14± 2.5%. While this reproducibility is lower than our previous studies using HeLa cells in a 5 mm probe (CV 2.4 ± 0.8%)^34^, it is important to note that we have established entirely new protocols for seeding, injection, and testing of the chip. The obtained CV corresponds well with typical values obtained in the literature^45^, where the CV across six studies ranged from 3% to 46%, with a mean CV of 23% ± 15% underscoring the impact of experimental conditions on reproducibility^45^. In this context, the current average CV of 13.66 ± 2.49% is significantly below the mean, indicating good reproducibility and measurement consistency. This performance is noteworthy given the complexity of integrating microfluidic systems with NMR technology. The chip is compatible with histological staining and optical microscopy, such as fluorescence and brightfield imaging (Figure 4), which is important for integrated use and benchmarking against other OOC platforms. Combining dDNP-NMR with fluorescence imaging provides complementary modalities, integrating functional metabolic measurements with static morphological insights.

### Comparative Analysis of One-Chamber and Two-Chamber Chips

Many OOC models utilize two-chamber chips, where the chambers are separated by a membrane onto which cells adhere^11^. This two-chamber design enhances physiological relevance and versatility, enabling more complex experimental setups and improving the functionality of OOC models. A key advantage of this configuration is the ability to implement differential flow between the top and bottom chambers, allowing cells in the top chamber to adhere undisturbed while nutrients are supplied through the bottom chamber. This is particularly important for growing epithelial monolayers of cells, which naturally form a barrier layer and need to be supplied with nutrients from the basolateral side of the monolayer to replicate physiological conditions^46,47^.

Using the same base design, we adapted our system for metabolic measurements in a two-chamber chip. This adaptation achieved similar cell densities as obtained in the one-chamber chip, with a confluent HeLa cell layer of approximately 4× 10^6^ cells per chip 24 hours after seeding (Figure 6A). To evaluate the system’s adaptability for different OOC configurations, we measured lactate production in the two-chamber chip. This design, commonly employed for barrier models, demonstrated metabolic activity comparable to the one-chamber chip. The two-chamber chip exhibited similar lactate production kinetics, with a calculated rate constant of 2.55 ± 0.28 × 10^−4^ s^−1^ (Figure 6B). These results highlight the robustness and flexibility of our system across different chip configurations, confirming its suitability for a wide range of biological applications, including barrier models and other complex OOC setups.

**Figure 6:**
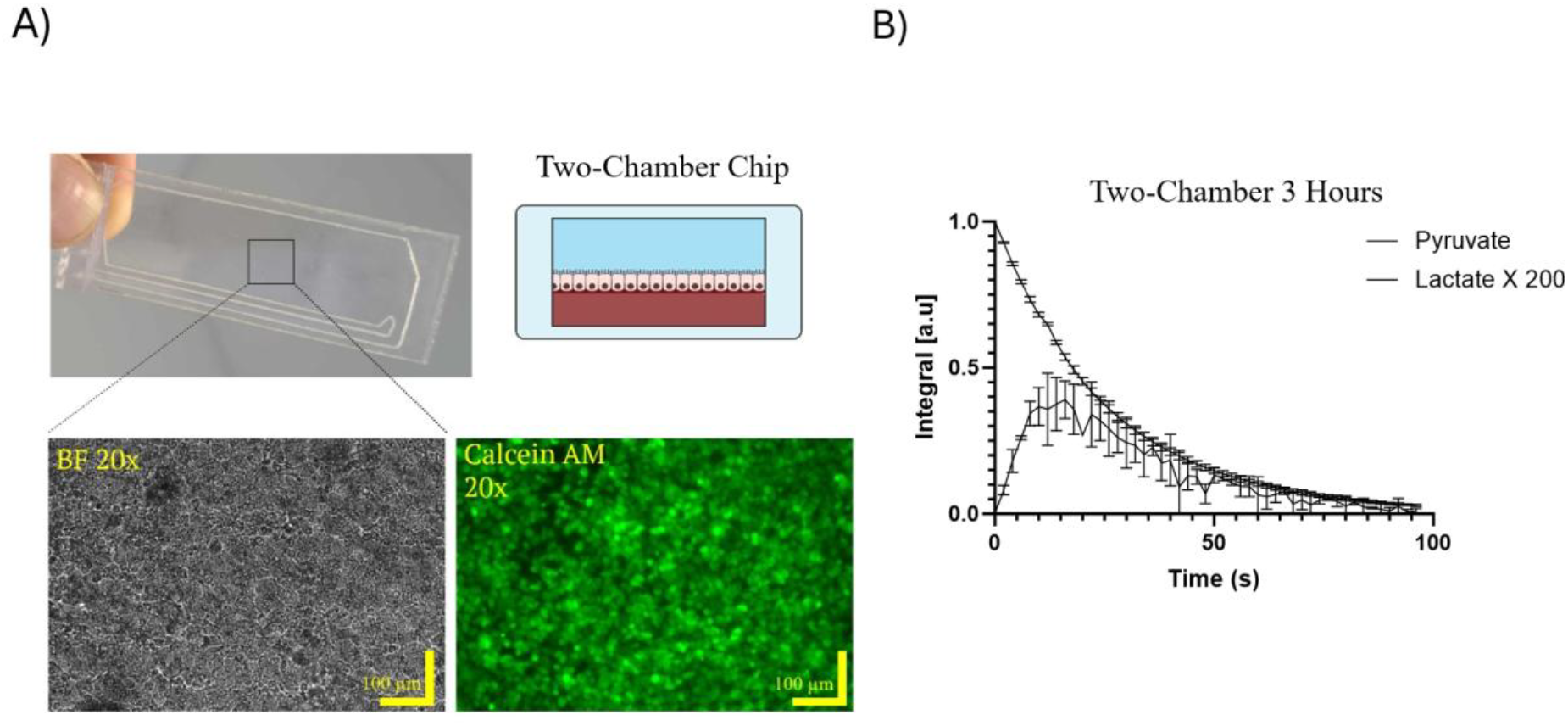
Two-Chamber Chip Cell Growth and Lactate Production: (A) Two-chamber chip seeded with 4.2 × 10^6^ HeLa cells, using a flow rate of 0.200 mL/min in the lower chamber, and no flow in the top chamber, for 24 hours at 37°C and 5% CO_2_. Bright-field microscopy image (bottom left, BF 20×) and fluorescence microscopy image of cells stained with Calcein AM (bottom right, Calcein AM 20×). Scale bar: 100 μm. The counted cell number was 3.03 ± 0.25 × 10^6^ (N = 3). (B) Time-series spectra showing [1-^13^C] lactate accumulation in the two-chamber chip seeded with 3.5 × 10^6^ HeLa cells, following injection of 10 mM hyperpolarized [1-^13^C] pyruvate. The mean ± SD rate constant for lactate production was 2.55 ± 0.28 × 10^−4^ (n = 2).

Our microfluidic chip successfully supports multi-day cell culture while maintaining stable metabolic function (Figure 4). Unlike conventional designs, which typically accommodate 10^3^–10^5^ cells, our chip can support up to 4 million cells in a monolayer^11,29^. As a proof of concept, we conducted metabolic assessments over 48 hours. Future studies should extend this timeframe to up to four weeks, which is necessary for more complex cocultures^48^. Long-term OOC cultures can develop into 4–8 cell layers, at full differentiation^49^. This would correspond to 12–36 million cells in the current chip. This increased cell density would significantly enhance sensitivity, possibly allowing for the detection of lower-concentration metabolic intermediates.

While the results are promising, several limitations and areas for future improvement should be noted. First, the current study focused on HeLa cells, a well-characterized cancer cell line. Future work should explore the system’s applicability to other cell types, including primary cells and co-cultures, to better replicate the complexity of human tissues and organs. The system is highly adaptable and can be extended to alternative cell models. For example, intestinal differentiation can be achieved in Caco-2 cells by asymmetrically culturing them with no fetal bovine serum (FBS) on the apical channel and with FBS on the basolateral side^50^. This protocol can be implemented on our two-chamber chip to analyze the metabolism of more complex and differentiated tissues, allowing for studies on epithelial barriers. Other cell types could also be used to model specific disease states or drug responses, further expanding the system’s applicability. The NMR probe demonstrated excellent signal stability, ensuring reproducibility across different experimental setups. Ensuring consistency across operators will be crucial for widespread adoption. Developing standardized protocols and calibration methods will be essential to address this challenge.

## Conclusion

Advanced cell models replicating features from human organs (OOC) have a high potential in reducing animal experiments and improving drug development processes. As the OOC technology continues to advance and demonstrate its value, the demand for OOC devices is expected to increase significantly in the near future^5^. Currently, options for non-invasive metabolic monitoring in OOCs are limited. Integrating metabolic measurements with OOC cellular models potentially enhances the physiological relevance, predictive power, and overall utility of these platforms. NMR, and in particular hyperpolarized NMR such as dDNP-NMR, offers real-time process monitoring in biological systems and integration of dDNP-NMR with microfluidic OOC technology marks a significant advancement in non-invasive, real-time metabolic analysis in these valuable cellular models.

This study presents a platform for non-invasive, real-time metabolic analysis in perfused microfluidic cell cultures. The integration of dDNP-NMR with custom-designed microfluidic chips enables dynamic monitoring of metabolic fluxes, offering a powerful tool for drug development, disease modeling, and personalized medicine. The system’s adaptability to different chip configurations and its ability to support long-term cell culture make it a versatile and valuable addition to the OOC toolkit. By bridging the gap between advanced NMR spectroscopy and microfluidic technology, this work paves the way for more accurate and human-relevant *in vitro* models, ultimately contributing to the development of safer and more effective therapies.

## Data Availability Statement

All data are available from the corresponding author upon request.

## Acknowledgements

This research was funded by the Danish National Research Foundation (grant DNRF124), EU Interreg Öresund-Kattegat-Skagerrak project ‘Hanseatic Life Science Research Infrastructure Consortium’ (HALRIC) and the Novo Nordisk Foundation (infrastructure grant NNF19OC0055825). JDSH contribution to this work was supported by Fundación Séneca (22401/SF/23).

