## Supporting Information for "ChipNMR: Hyperpolarized NMR for non-invasive metabolic flux analysis in perfused microfluidic chips"

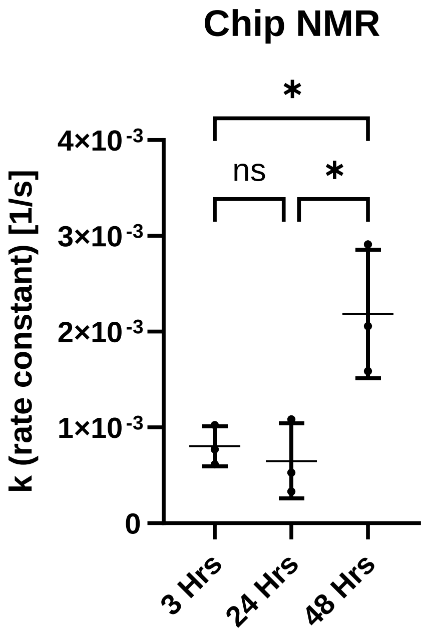


**Figure SI 1:** Rate constants from dDNP NMR measurements in microfluidic chips following injection of 10 mM hyperpolarized [1-$^{13}$C] pyruvate. Rate constants (mean ± SD) were 0.000804 ± 0.000170 at 3 h, 0.000649 ± 0.000319 at 24 h, and 0.002185 ± 0.000548 at 48 h. Corresponding cell counts (mean ± SD, millions) were 1.90 ± 0.26 at 3 h, 2.70 ± 0.61 at 24 h, and 4.46 ± 0.69 at 48 h.
